# mRNA and Small RNA-seq Reveal Insights into Immune Regulation in *Apis cerana* after Chinese Sacbrood Virus Infection

**DOI:** 10.1101/2020.07.07.192799

**Authors:** Yanchun Deng, Hongxia Zhao, Shuo Shen, Sa Yang, Dahe Yang, Shuai Deng, Chunsheng Hou

## Abstract

Chinese sacbrood virus (CSBV) is a serious threat to eastern honeybees (*Apis cerana*), especially larvae. However, the pathological mechanism of this deadly disease is remains unclear. Here, we employed an mRNA-seq and sRNA-seq approach in healthy and CSBV-infected 3^rd^ *Apis cerana* larval. Gene ontology (GO) and KEGG analysis of 203 differentially expressed genes showed that CSBV infection affected host development by up-regulating the expression of larval cuticle proteins, such as larval cuticle proteins A1A and A3A, resulting in elevated susceptibility to CSBV. In addition, viral infection not only affected the expression of serine protease related to the melanization pathway and but also altered fatty acid metabolism and biosynthesis, thus progressed to disturb host immune response. Interestingly, GO annotation and KEGG analysis on target genes of CSBV-specific siRNA (vsiRNAs) showed that serine/threonine kinase activity and serine-type endopeptidas as well as fatty acid biosynthesis were significantly enriched (P < 0.05). Among these vsiRNAs, vsiRNA-1441 with relatively high expression targeted extracellular serine/threonine protein kinase. This study provides new evidence that CSBV attacks a distinct immune response pathway and mediates the expression of cuticle protein to gain the more chance of proliferation.

**IMPORTANCE:** Chinese sacbrood virus (CSBV) is the major factor threatening the health of *Apis cerana*. Although honeybees have antiviral defense mechanisms including RNA-interference (RNAi), virus had evolved resistance strategy like virus-derived small interfering RNAs (vsiRNA) produced during the viral infection, which can alter host immune by hijacking the host RNAi pathway. To gain insights at the transcription level into the mechanisms vsiRNA altering immune response and how this variation is connected with CSBV infection, we performed the first combined mRNA and small RNA-seq analyses of healthy and CSBV-infected *Apis cerana* larvae. Together, the omics approach enabled us to obtain comprehensive results that CSBV infection changed the expressions of cuticle proteins and related serine protease, and leading to the larvae unable to pupate. This will guide the design of new strategies for control CSBV.

---

The Asiatic honeybee, *Apis cerana*, is an important pollinator to maintain plant biodiversity in Southeast Asian countries, especially on wild flowering plants and crops **(1)**. However, honeybees have been suffering from colony decline in recent years due to an increasing number of infections from multiple pathogens **(2)**. Among honeybee pathogens, Chinese sacbrood virus (CSBV), a Chinese strain of sacbrood virus (SBV) obtained from *A. cerana*, is the major threatening factor in *A. cerana* colony decline **(3)**. SBV, as the only single-strand positive RNA virus, can infect honeybee larvae and lead to larval death **(4)**. SBV belongs to the genus *Iflavirus* and one of members of the family *Iflaviridae* within the order *Picornavirales*, which includes several important honeybee viruses, such as deformed wing virus **(5)**. The full-genome sequence of SBV was acquired in 1999 **(6)**.

SBV-infected *A. cerana* was termed CSBV. The clinical symptom of infected larvae was ecdysial fluid aggregated with typical “sac”, resulting in failure to pupate **(7)**. CSBV, obtained firstly from bee samples in Guangdong province of China in 1972 **(8)**, has frequently caused extensive larvae death and colony decline and recently re-emerged as one of the causative agents of larva disease outbreaks in 2008 in Liaoning **(9)**. Since then, the virus has frequently infected *A. cerana* in this region of China **(10)** and remains a major threat to *A. cerana* colonies in China **(3)**. Usually, the viral infection can induce cell apoptosis and tissue damage as well as functional disorder. Then, all these alterations can be seen through gene expression changes **(11, 12)**.

Like other insects, honeybees have no adaptive immune response known in vertebrates **(13)**. To survive under the persistent threat of viral infection, honeybees have evolved several defense mechanisms that are activated via different immune pathways **(14, 15)**. Honeybee antiviral defense mechanisms include RNA-interference (RNAi), Janus kinase/Signal Transducer and Activator of Transcription (Jak/STAT) pathway, Toll pathway, Immune Deficiency (Imd) pathway, c-Jun N-terminal kinase (JNK) pathway, Mitogen-Activated Protein Kinases (MAPK) pathway and melanization, as well as reactive oxygen species generation **(16)**. Serine protease/proteinase (SP) and serine protease/proteinase homolog (SPH) play an important role in innate immune response that includes coagulation, melanisation and the Toll signaling pathway in honeybee **(15, 17)**.

Experimental evidence by quantitative PCR (qPCR) has shown that CSBV infection induces routine immune responses, such as activating the expression of antimicrobial peptides **(18)**. Generally, activation of the Toll, Imd and Jak/STAT pathways result in the production of antimicrobial peptides and other effector proteins **(16)**, nevertheless, knowledge about the comprehensive immune responses to CSBV infection is limited. Omics technique is a useful tool to measure the dynamic changes of whole gene expression associated to viral infection **(19)**. Using the proteomic technique, it was found that there were 142 proteins and 12 phosphoproteins downregulated in CSBV-infected larvae, the infected worker larvae were significantly affected by such as carbohydrate and energy metabolism and fatty acid metabolism **(20)**. Lately, the full-genome sequence of *A. cerana* from Chinese honeybee was determined, providing new insights into physiological resistance to *Varroa* mites, and was found to have 6 more immune genes than *A. mellifera* **(21)**. In addition, transcriptome technology was applied to several study fields in honeybees, such as gland development **(22)**, *Varroa* mite control **(23)**, and *Ascosphaera apis* pathogenesis **(24)**. Previous studies have found that challenges by mechanical stressors and pathogens can lead to significant immune responses in honeybees, including a significant upregulation of *abaecin, defensin1, apidaecin*, and *hymenoptaecin* genes **(25, 26)**. Galbraith et al. (2015) identified the transcript and epigenetic responses of honeybees to Israeli acute paralysis virus (IAPV) infection and found that honeybees employed distinct different immune pathways, such as JAK-STAT, in resistance to viral infection apart from universal immune responses **(27)**. Recently, Rutter et al. **(28)** confirmed that IAPV infection combined with diet quality can impact the immune gene expression of Notch and JAK-STAT signaling pathways. Nevertheless, artificial infection with acute bee paralysis virus (ABPV) did not induce a humoral immune response of bee larvae and workers because artificial infection frequently induced the unexpected transcription changes of antimicrobial peptides genes **(29)**. For example, artificial infection or injection to bee larvae as an experimental control results in changes of transcription patterns in non-target genes **(30)**.

Small RNAs include microRNAs (miRNAs) and small interfering RNAs (siRNAs) that are involved in regulating gene expression in most organisms **(31, 32, 33)**. Virus-derived small interfering RNAs (vsiRNA), are a group of siRNA in size ranged from 21-24 nt that were produced during the viral infection, have several functions such as a type of siRNA that may hijack the host RNAi pathway **(34)**. In brief, vsiRNA, which guides the RNA induced silencing complex (RISC) to target viral genomes in invertebrates **(33)**, are associated with the RNAi-based antiviral response can potentially alter the host transcriptome responses **(35)**. During viral replication, most viruses produce long segments of dsRNA, including replication intermediates of ssRNA viruses and RNA secondary structures in long transcripts of viruses. Long exogenous dsRNAs are cleaved by Dicer into siRNA duplexes of 21 to 22 nt **(36)**. Endogenous miRNAs are canonically derived from transcribed miRNA precursors encoded in protein coding or non-coding transcription units in the genome by RNA polymerase II. miRNAs are a group of sRNAs with size 22 nt and are significant regulators of diverse biological processes, including development and interactions between host and virus **(33)**.

Thus, the mechanisms underlying host responses to CSBV infection are unknown, especially the transcriptomical level and the regulatory effects of small RNAs to CSBV infection under natural conditions. Here, we examined the transcriptional responses and the abundance of siRNAs of *A. cerana* larvae to CSBV infection under natural conditions. We characterized (1) the transcriptomic expression of cuticle proteins related to larval development and immune-related genes, (2) host metabolism, and (3) determined whether there is a relationship between siRNA and CSBV infection. Our results provide insights into up-regulated cuticle protein genes and down-regulated serine proteinase and fatty acid biosynthesis to facilitate the viral infection. These findings significantly broaden our knowledge of virus-host interactions and provide novel targets for the control of CSBV.

## RESULTS

### The overview of expressed mRNA and small RNAs

A total of 6 RNA-seq libraries were obtained using Illumina sequencing platforms, but we subjected 2 pooled controls and 2 pooled CSBV infected samples to further analysis due to the 2 poor data with other virus contamination (data not show). For sRNA sample sequencing, a total of 18,270,587 (T01), 19,574,141 (T02), 17,646,189 (T03) and 20,618,929 (T04) clean reads were obtained from the healthy and CSBV-infected mRNA libraries (Table S1) (T01 and T02 belong to the healthy blank groups; T03 and T04 belong to the CSBV-infected sick group). Mapped reads to *A. cerana* genome were more than 70% among all samples (Table S2). Meanwhile, for sRNA sample sequencing, a total of 18731778 (T01), 17407255 (T02), 15937441 (T03) and 13237265 (T04) clean reads were obtained from the healthy and CSBV-infected sRNA libraries (Table S3). The value of Q30 was more than 90% among all samples, which indicated that these clean reads can be used for subsequent analysis (Table S1 and S3).

We further studied the expression levels of all mRNAs between control and CSBV-infected larvae based on FPKM value (fragments per kilobase of exon per million fragments mapped FPKM) (**Fig. 1A**), and then found that the FPKM value of mRNAs distributed from 0 to 10000 in the four libraries, but the mean FPKM value in T01 is slightly lower than the other three libraries (**Fig. 1A)**. The correlation analysis on two pooled group using R.ggplot 2 confirmed that it is unreasonable to make T01 and T02 together in healthy group (Correlation=0.275) and reliable to make T03 and T04 together in CSBV-infected group (Correlation=0.622) (**Fig. 1B and 1C**).

**FIG 1.**
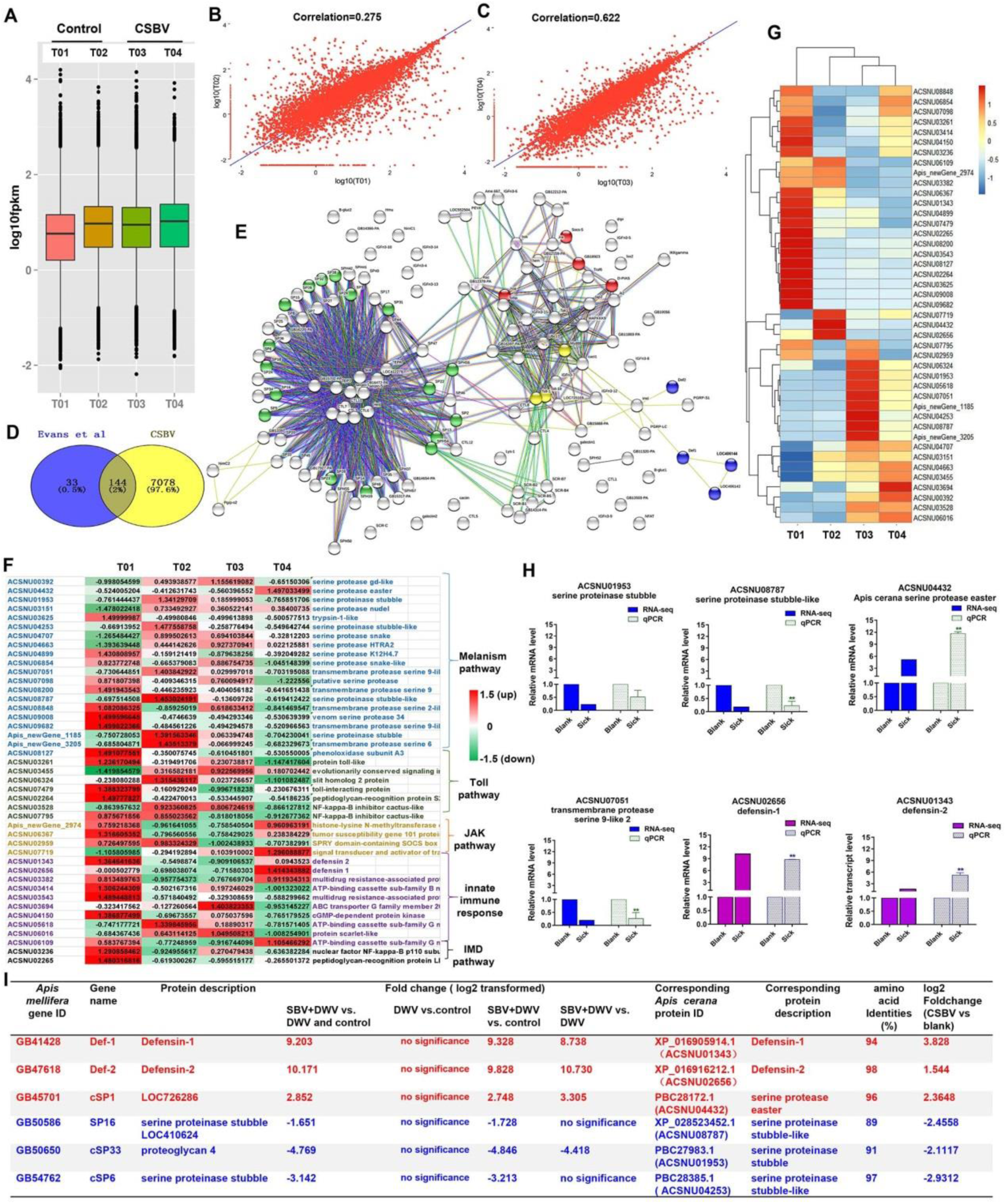
The correlation analysis of all samples. (A) The analysis of expression level of samples based on FPKM value, T01 and T02 belong to group, T03 and T04 belong to CSBV-infected group. (B) Correlation analysis of the expression of all mRNA between healthy duplicated samples. (C) Correlation analysis of the expression of all mRNA between CSBV-infected duplicated samples. (D) One hundred forty-four expressed immune genes were shared between this study and described by Evans et al. (2006). (E) The predicted interaction network of 144 shared genes, and 4 major terms were enriched (P<0.05). Green, serine proteases and trypsin domain; red, Jak-STAT signaling pathway; blue, innate immune response; yellow, Toll receptor homology domain. (F) The Statistic analysis of the regulated genes related to serine proteases, melanization and Toll pathway, Jak-STAT signaling pathway, innate immune response, and IMD pathway in honeybee larvae after CSBV infection. The values represent standardized expression levels based on FPKM mean values. Green and red indicate decreased and increased in expression levels, respectively. (G) Heat map of the regulated genes related to serine proteases, melanization and Toll pathway, Jak-STAT signaling pathway, innate immune response, and IMD pathway in honeybee larvae after CSBV infection. (H) The relative expression levels of serine proteinase and defensin genes from RNA-seq (T02, T03 and T04) and qPCR analyses. Actin was used as the internal reference gene. *P<0.05,**P<0.01. (D) The differentially expressed serine protease and defensin genes (P<0.05, FDR <0.05) in our study were compared with that of SBV-induced (Ryabov et al., 2016) (P<0.05, FDR <0.05).

### Comparative analysis of immune genes against CSBV infection with other studies

To detect the difference in immune response among healthy and CSBV-infected honeybee samples, comparative analysis of immune genes involving in against CSBV infections with the other study was performed. A comparison of our suite of 7,222 regulated genes (FDR <0.05 and P <0.05) and the 177 genes related to honeybee innate immune system described by Evans et al. **(13)** showed they shared 144 genes (**Fig. 1D**), which was consistent with those of *A. cerana* immune response **(11)**. Then, we predicted the interaction networks of 144 genes and found 17 serine proteases genes (P <0.0001) highly related to melanization pathway, 4 genes related to the JAK-STAT signaling pathway (P <0.01), 3 Toll receptor homology domain genes (P <0.05), and 4 genes related to innate immune responses (P <0.001), respectively **(Fig. 1E)**. The statistical analysis of standardized expression levels of the regulated genes related to serine proteases, melanization and the Toll pathway, Jak-STAT signaling pathway, innate immune response, and IMD pathway in honeybee larvae after CSBV infection showed that a majority of up-regulated immune genes such as *defensin* were identified in T01 samples (**Fig. 1F**). Clustering heat map analysis of these genes confirmed that T02 was health group and T3 with T4 were infected group, respectively (**Fig. 1G**). In addition, it is difficult to find healthy samples under natural conditions **(11, 37)**. Therefore, T02 as the control, T03 and T04 as CSBV-infected group were used to analyze the relative expression levels of serine proteinase and *defensin* genes (**Fig. 1H**). Based on FPKM value and Ct value of immune genes, *defensing* 1 and *defensing* 2 were significantly up-regulated **(Fig. 1H)**. The transcriptional levels of serine protease genes were variable. For example, serine proteinase stubble, serine proteinase stubble-like and one transmembrane protease serine 9-like were downregulated to 0.5-, 0.22- and 0.26-fold, while serine protease easter had > 11.68-fold elevated expression (**Fig. 1H**), which was consistent with RNA-seq analysis (r =0.8004, P =0.05)(**Fig. S1**).

Next, we performed comparative analysis of main immune responses to CSBV infections with the previous SBV study **(38)**. Ryabov et al. **(38)** found that worker pupae orally infected with SBV and deformed wing virus (DWV) resulted in down-regulation of genes involved in cuticle and immune compared to orally infect with only DWV. In our study, several differentially expressed *defensin* and serine protease genes were consistent with those of SBV induced (only P <0.05, FDR <0.05 and NCBI blast amino acid identities >80% were listed) **(Fig. 1H)**. For example, serine proteinase stubble-like and serine proteinase stubble genes were significantly down-regulated (P <0.05), while *defensin* 1, *defensin* 2 and serine protease easter were up-regulated **(Fig. 1H)**. Additionally, qPCR results showed that the expression of the *PPO* (*prophenoloxidase*) gene was significantly down-regulated at the 4^th^, 5^th^, 6^th^, and 7^th^ instar after CSBV infection (**Fig. S2**).

### Gene ontology (GO) and KEGG analysis on differentially expressed mRNA involved in response to CSBV infection

A total of 10,262 mRNA transcripts were produced (FDR <0.05) **(Table S4)**, and 203 significantly differently expressed genes (DEGs) (**Table S5**, log2 FoldChange >1.5 or < -1.5, Probability > 0.8) were identified. Among the 203 DEGs, 83 genes were successfully matched to GO terms **(Fig. 2A)**, and the top 10 GO terms (P ≤0.01) of molecular function belong to the structural constituent of cuticle, serine-type endopeptidase, serine-type peptidase and serine hydrolase activity (**Fig. 2A)**. Based on GO-directed acyclic graph the relationship among these 10 terms in molecular function showed that the structural constituent of cuticle and serine-type endopeptidase activity were significantly enriched (P <0.01) (**Fig. 2B)**. Ten genes related to the structural constituent of cuticle were significantly up-regulated including Larval cuticle protein A1A, larval cuticle protein A2B, flexible cuticle protein 12-like, and larval cuticle protein A3A **(Table S5)**. While 7 serine protease genes were significantly down-regulated except easter was remarkably up-regulated.

**FIG 2.**
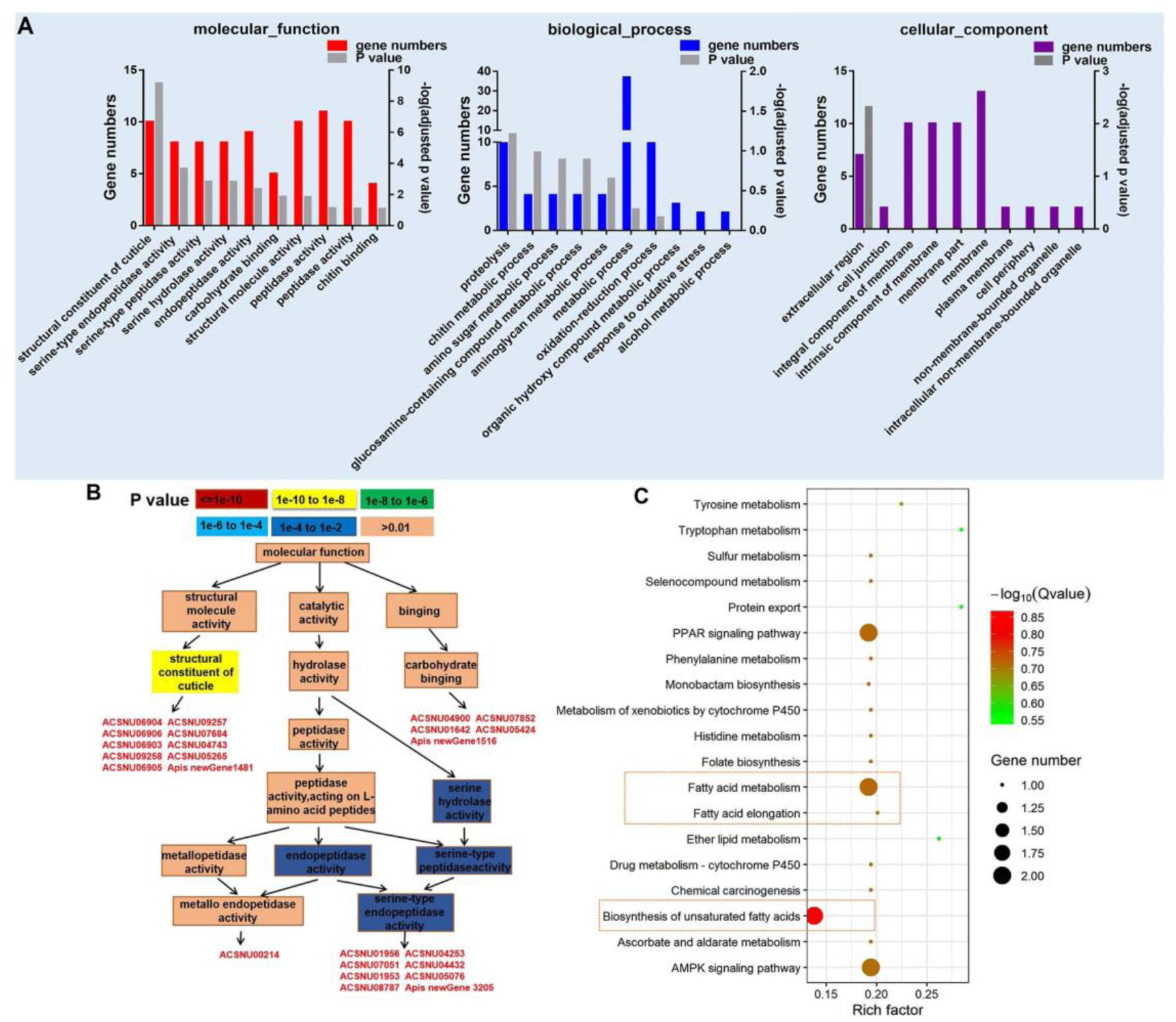
Gene ontology (GO) analysis and KEGG analysis differentially expressed genes (DEGs). (A) The first 10 GO terms analysis (P < 0.05) of DEGs in molecular function, biological process and cellular component. (B) GO-directed acyclic graph chart which belongs to molecular function. The color represents the enrichment significance levels according to *P*-value. (C) The KEGG analysis of DEGs (P < 0.05).

KEGG analysis on 203 DEGs found that fatty acid metabolism, biosynthesis of unsaturated fatty acids, AMPK (Adenosine 5’-monophosphate (AMP)-activated protein kinase) signaling pathway, and PPAR (Peroxisome proliferator-activated receptors) signaling pathway were mainly enriched in 20 categories **(Fig. 2C)**. Interestingly, two acyl-CoA Delta 11-desaturase-like genes (ACSNU05780 and ACSNU05780) involving in fatty acid metabolism, biosynthesis of unsaturated fatty acids, AMPK signaling pathway, and PPAR signaling pathway were significantly up-regulated to 4 fold. Furthermore, we also found that two long chain fatty acids protein genes related to fatty acid biosynthesis among 203 DEGs were significantly down-regulated to 0.12 fold at least **(Table S5)**.

### GO analysis and KEGG annotation on target genes of differentially expressed miRNA involved in response to CSBV infection

Using miRDeep2 and DESeq software, we mapped clean sRNA reads to the *A. cerana* genome against the miRanda database **(39)**. The compositions of those reads of sRNA are shown in **Fig. S3**, and the most abundant class of sRNAs were 22-nt in size, including known miRNA and novel miRNA (**Fig. S3A and B)**. Two hundred sixty miRNAs were obtained, which included 23 differently expressed miRNAs (**Table S6**, log2 FoldChange > 0.6 or <-0.6, P < 0.05). Using bioinformatic prediction method, a total of 1035 target genes were identified by 260 miRNAs and 23 differently expressed miRNAs were predicated towards 227 target genes that involved in biological process, cellular component, and molecular function, and most of them were identified to belong to reproduction, nuclear chromosome and nucleotide binding, respectively (**Fig. 3A**). KEGG annotation analysis showed that 227 target genes mainly participate in ECM-receptor interaction, ribosome biogenesis in eukaryotes and endocytosis (**Fig. 3B**).

To examine whether the results of miRNA were consistent with those of mRNA, we identified miRNAs and found the expression of ame-miR-3759 was significantly up-regulated to more than 1-fold (**Table 1**). While the expression of the 2 targeted genes, LOC410515 and putative uncharacterized protein DDB_G0277255-like, were significantly down-regulated (P <0.01). LOC410515 may have a similar role in immune function with homologous protein *Apis dorsata* chorion peroxidase-like (XP_006611124.1) (NCBI blast sequence identify >75%), which played an important role in melanin synthesis **(40)**.

**TABLE 1.** The list for opposite expression changes in miRNA-mRNA pairs

| miRNA | miRNA_log2<br>Foldchange | Target<br>mRNA | mRNA_log2<br>Foldchange | Annotation | Homologous<br>protein |
| --- | --- | --- | --- | --- | --- |
| ame-miR<br>-3759 | 1.09 | ACSNU0<br>5809 | -3.40 | putative<br>uncharacterized protein<br>DDB_G0277255-like | peptidyl-prol<br>yl cis-trans<br>isomerase |
| ame-miR<br>-3759 | 1.09 | ACSNU0<br>5034 | -3.61 | uncharacterized<br>protein LOC410515 | chorion<br>peroxidase |

**FIG 3.**
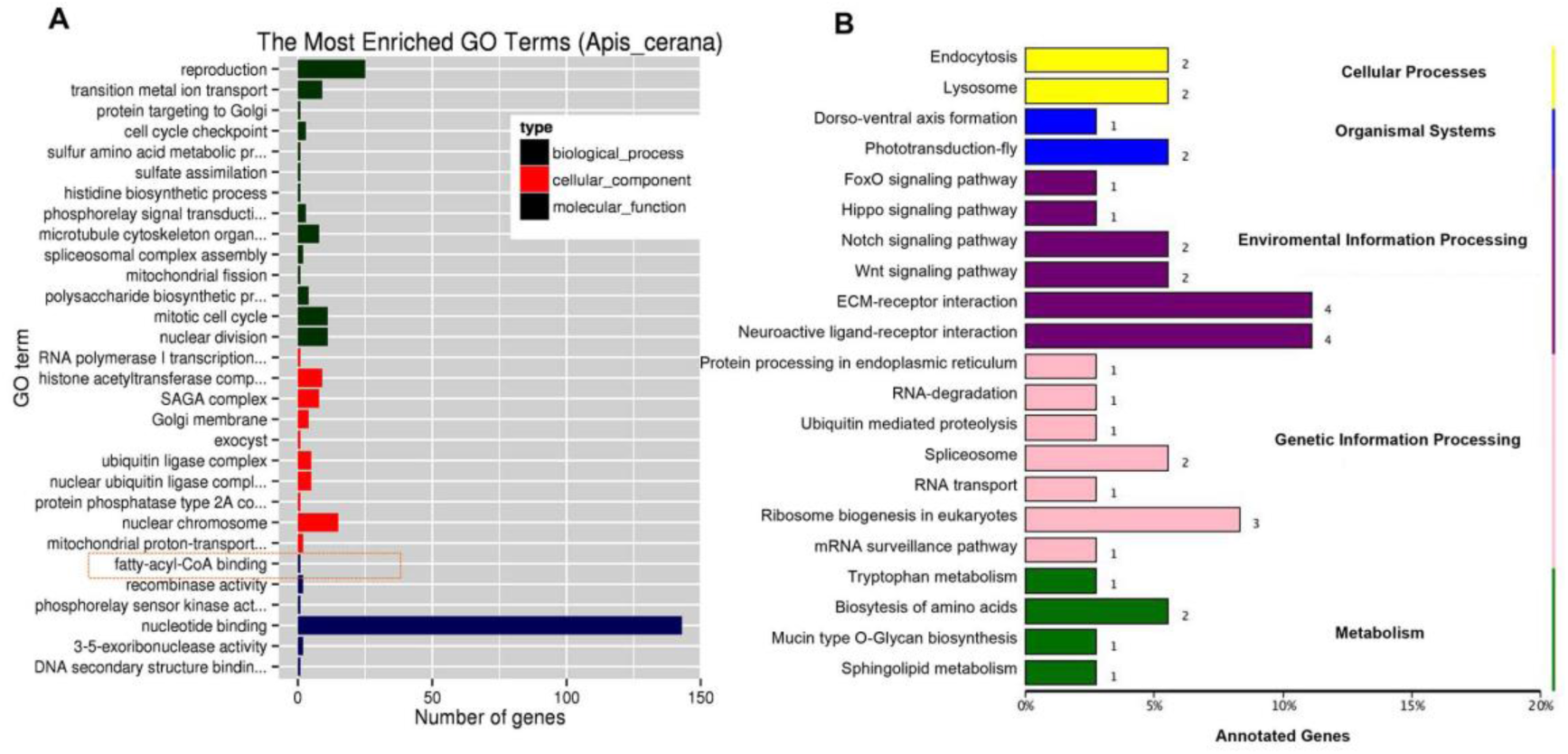
GO analysis and KEGG annotation on target genes of differentially expressed miRNA. (A) The most enriched GO terms (P < 0.05) of target genes of differentially expressed miRNA in molecular function, biological process and cellular component. (B) The KEGG analysis of target genes of differentially expressed miRNA (P < 0.05).

### Identification, GO annotation and KEGG analysis on target genes of CSBV-specific siRNA

We aligned all the cleaned sRNA-seq reads to the CSBV genome (GenBank: KU574662.1) using a known miRNA-target database (miRanda, 3.3a) and 2,467 vsiRNAs were obtained, in which 319 effective vsiRNAs could be predicted to a total of 290 target genes (**Table S7, Table S8**), which were mainly related to DNA binding, transcription factor activity, apoptosis, G-protein coupled receptor activity, transporter activity and serine/threonine kinase activity by molecular functions prediction (**Fig. 4A, Table S8**). KEGG analysis on target genes of CSBV-specific siRNA showed that purine metabolism, hippo signaling pathway and fatty acid biosynthesis were significantly enriched (P < 0.05) (**Fig. 4B**).

**FIG 4.**
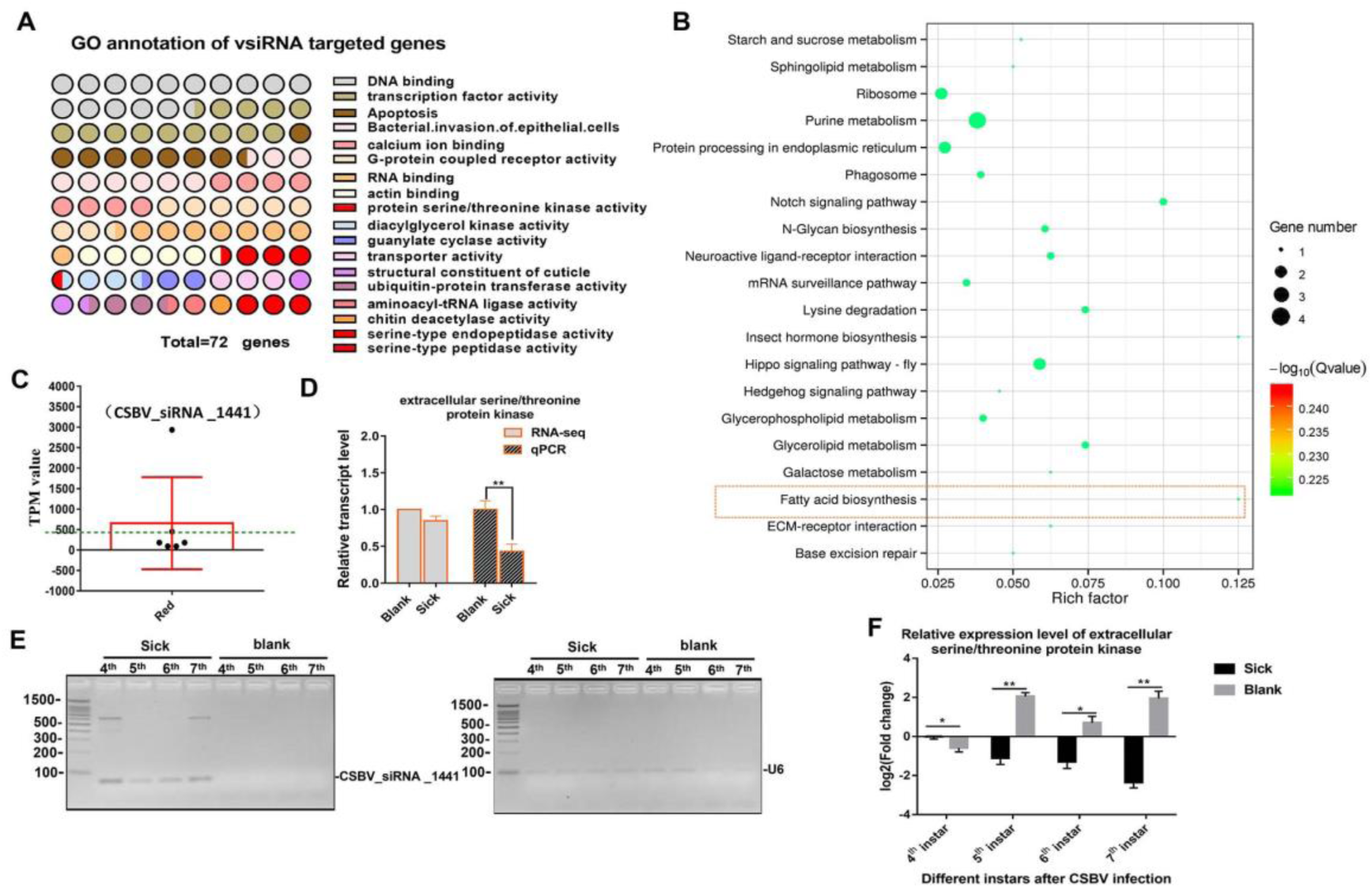
GO and KEGG annotation of vsiRNAs target genes. (A) GO annotation of vsiRNAs target genes in CSBV infected larvae of Asian honeybees. (B) The KEGG analysis of vsiRNAs target genes. (C) The target gene of vsiRNA_1441 was extracellular serine/threonine protein kinase FAM20C-like of Apis (the TPK value was higher than mean value). (D) The relative expression level of the serine/threonine protein kinase FAM20C obtained from RNA-seq and qPCR analyses. (E) Analysis of miRNAs in 3% gel displaying continuous expression in CSBV-infected honeybees at different instars. U6 RNA is used as a loading control. (F) qPCR analyses of the relative expression levels of serine/threonine protein kinase FAM20C in CSBV-infected honeybees at different instars. *P<0.05,**P<0.01.

Although the expression of dicer-like and argonaute-2 (Ago2) genes associated to the RNAi pathway were up-regulated (**Table S3, Fig. S4**), we focused on siRNAs related to serine/threonine kinase, serine-type endopeptidase and serine-type peptidase genes based on the analysis of mRNA and miRNA data. Then, the expression analysis of vsiRNAs related to serine/threonine kinase, serine-type endopeptidase and serine-type peptidase were performed based on the TPM (Transcripts Per Million Reads) value of siRNAs (**Fig. 4C, Table S9**). Generally, the complete expression profile of vsiRNAs was categorized into ‘low’ (<50 TPM value), ‘high’ (>=50 TPM value) and ‘very high expression’ (>=1000 TPM value) (relative abundance) **(41)**. The expression levels of vsiRNA_1441 (TPM value is 2932) were extremely significant higher than other siRNAs (3 to 20 times higher, mean value is 405) (**Fig. 4C**). Then, we found that the target gene of vsiRNA_1441 was extracellular serine/threonine protein kinase FAM20C-like of *Apis*, which potentially regulate many processes including proliferation, cell survival, growth, angiogenesis and metabolism. Subsequently, qPCR analysis confirmed that the expression of extracellular serine/threonine protein kinase FAM20C was significantly down-regulated (**Fig. 4D**). Next, the expressions of vsiRNA_1441 at different instars of CSBV-infected larvae were validated by RT-PCR (reverse transcription PCR) (**Fig. 4E**). Similarly, the expression of the extracellular serine/threonine protein kinase FAM20C gene was significantly down-regulated at the 4^th^, 5^th^, 6^th^ and 7^th^ instar after CSBV infection (**Fig. 4F**).

## DISCUSSION

Healthy larvae are crucial for the development and growth of the *A. cerana* colony population. However, CSBV contributed to recent declines of the *A. cerana* population **(11)**. A few studies have shown that vsiRNAs, as pathogenicity determinants, negatively regulate host mRNAs and effectively silence host genes to gain more proliferation **(35)**. Here, to understand the underlying molecular pathological mechanism of the CSBV infection, we presented the first transcriptome and siRNA analysis from CSBV under natural conditions. Accordingly, results of RNA-seq showed that 203 genes were significantly altered between the healthy and infected larvae. Of these, all cuticle protein genes related to structural constituent of cuticle were significantly up-regulated, while serine protease genes related to serine-type endopeptidase activity, serine-type peptidase activity and serine hydrolase activity were significantly down-regulated. Meanwhile, 23 differently expressed miRNAs and 319 effective vsiRNAs were identified, which some of vsiRNAs targeted serine/threonine kinase or serine-type endopeptidase-related genes. These findings provide a novel insight into how CSBV resists host immune responses and prevents larvae from reaching to the pupae stage.

### Cuticle proteins play a key role in CSBV infection

Cuticle is a barrier against viral invasion in animals such as white spot syndrome virus infection in shrimp **(42)**. Cuticle proteins play an critical role in keeping the body from pathogens and serving as a barrier to resistance **(43)**. It has been reported that natural resistance-associated macrophage protein (NRAMP) was a cellular receptor of Sindbis virus in insects **(44)**. In our study, NRAMP was not enriched by analyzing the DEGs. In *aphidiae*, cuticle protein 4 is essential to aphid acquisition of cucumber mosaic virus (CMV) because it was considered as a viral putative receptor **(45)**. While in honeybee, apidermin 3, a cuticular protein was down-regulated after alone deformed wing virus infection but up-regulated after deformed wing virus and *Varroa* mite infection **(46)**. Our results showed that the expression levels of larvae cuticle protein genes including larval cuticle protein A2B, larval cuticle protein A1A, larval cuticle protein A3A were greatly elevated, while Ryabov et al. **(38)** found that the genes involved in cuticle and muscle development were down-regulated at a later stage after SBV infection (9 dpi). The transcription expression of some cuticle proteins was reported to be significantly up-regulated or down-regulated during different stages of the white spot syndrome virus **(47)**. Thus, we infer that larvae structural constituent of the cuticle may play an important role in CSBV infection.

### Down-regulated serine proteases play a vital role in suppression of the melanisation pathway in the CSBV-infected larvae

Serine proteases (SPs) with 60-400 members in insects form a large family of enzymes that hydrolyze peptide bonds at different rates **(48, 49)**. SPs, especially extracellular SPs, have a great influence on insect development and innate immunity **(50)**. SPs and SP homologs (SPHs) are involved in multiple physiological processes of insects such as digestion, defense and development **(48)**. Experimental evidence showed that SPs of *A. mellifera* may be involved in the embryonic development and immune responses **(15)**. Moreover, SPs are fundamental to melanization and the Toll pathway signaling **(51)**. Our results suggested that most immunity genes that overlapped with Evans’ study were related to the SPs. In this study, RNA-seq analysis showed that SPs related to serine-type endopeptidase and peptidase activity were significantly down-regulated. qPCR result also confirmed that CSBV infection lead to down-regulation of SP genes, such as SP stubble and transmembrane protease serine 9, and this was consistent with that of sacbrood virus infection **(38)**.

It also has reported that the viral proteins encoded by SBV may directly and simultaneously activate the Toll and Imd pathways, and the expression of *defensin*-1 and *defensin*-2 was significantly up-regulated in orally infected larvae with high dose of SBV **(38)**. These results showed that CSBV strains from *A. cerana* induced similar immune responses to those of SBV in *A. mellifera*. Honeybees can initiate humoral immunity involving SPs that include coagulation, melanization, and the Toll signal pathway **(52)**. SPs exist in hemolymph in the form of zymogens and then activate downstream proteins. Although Gao et al. **(17)** identified an SP gene, *Accsp10*, related to development and pathogens resistance from *A. cerana*, they did not confirm its function. However, our results showed that the *ppo* gene in the melanization pathway were down-regulated. The observed suppression of the melanization pathway in the SBV-infected larvae resulted from down-regulated prophenoloxidase-activating enzyme **(38)**. Additionally, we also found that the expression level of the *ppo* gene was significantly down-regulated at the 4^th^, 6^th^, and 7^th^ instar in CSBV-infected larvae and suggested that melanization involved in the resistance to CSBV infection were significantly suppressed. This is similar to that of the suppression of Semliki forest virus on the phenoloxidase cascade in mosquito **(52)**. CSBV infection inhibited the immune response of serine proteases, while intermediate steps of the cascade are still unknown.

### vsiRNAs regulated the expression of immune genes and viral infection disrupt host metabolism

Small RNAs (sRNA) have been confirmed to be related to a diversity of biological processes including cell proliferation and apoptosis through the post-transcriptional regulation of gene expression **(53-55)**. RNA profile analyses suggested that these sRNA are involved in biological processes associated with development and immunity of honeybees **(56)**. It has also been reported that vsiRNAs can regulate the gene expression related to the host RNAi pathway and enhance the pathogenesis. For example, Xu et al. **(57)** found that siRNAs derived from Southern rice black-streaked dwarf virus targeted several types of genes related to host resistance, such as receptor-like protein kinases. Although Chejanovskey et al. (2014) identified the viral siRNA population from colony with colony collapse disorder, they had no further analysis on targeted genes of siRNA. Recently, Chen et al. **(58)** feed siRNA target the viral suppressor of RNAi and significantly suppressed IAPV replication. The key initiator of the RNAi pathway is double-stranded RNA (dsRNA), which produced by proliferation process are processed into vsiRNA duplexes by dicer-2 **(59)**. Viral small interfering RNAs were processed into 21-nt through Dicer-2 on viral dsRNA and then are integrated into insect Ago2 and guide the Ago2 onto target RNAs to cause their degradation **(59)**. In our study, the expression of dicer-like and Ago2 genes were up-regulated and suggested that various vsiRNA could affect the expression of host genes by RNAi pathway. For instance, the predicted a total of 290 target genes of the vsiRNAs mainly related to DNA binding, transcription factor activity, apoptosis, G-protein coupled receptor activity, transporter activity, serine/threonine kinase activity and serine-type endopeptidase. Of these, vsiRNA_1441 significantly suppressed the expression of extracellular serine/threonine protein kinase FAM20C-like. Several studies have reported that serine/threonine kinases (STKs) are essential to regulate various cellular processes, metabolism and stress responses **(60, 61)**.

Previous studies have mainly investigated host response to viral infection. However, metabolites are another important indicator that can reflect physiological response to pathogens **(62)**. Fatty acid metabolism and biosynthesis is known to play a vital role in viral infections and proliferation in animals **(63, 64)**. It has reported that fatty acids can influence host immune by affecting the inflammatory repertoire of the host, substrates for biosynthesis of inflammatory mediators and activation of cell receptors **(65)**. In our study, KEGG analysis showed that 203 DEGs were significantly enriched in fatty acid metabolism and biosynthesis which was found in integrated KEGG analysis of DEGs and vsiRNA target genes. Meanwhile, two acyl-CoA Delta 11-desaturase-like genes related to fatty acid metabolism, biosynthesis of unsaturated fatty acids, AMPK signaling pathway and PPAR signaling pathway were significantly up-regulated after CSBV infection, while two long chain fatty acids protein genes related to fatty acid biosynthesis were significantly down-regulated. PPAR signaling pathway is considered as one of critical regulators in fatty acids metabolism and immune system **(66)**. Siwen et al. found that the AMPK activity was positively related to PRRSV (porcine reproductive and respiratory syndrome virus) infection and result in a decline of acetyl-CoA carboxylase 1 (ACC1) activity **(67)**. In addition, fatty acids were used to control the American foulbrood and other bee diseases of honeybee **(68)**. These evidences suggested that roles of fatty acid metabolism and biosynthesis in immune response could be critical factors that regulate physiological resistance to CSBV. However, further studies are needed to clarify the detail process.

In a world, according to result from combination previous study with this study, thus, we propose a hypothesis that CSBV deploy vsiRNA to modulate honeybee immunity. During early stages of infection, CSBV enter into the honeybee cells and host could up-regulate RNAi pathway to against CSBV proliferation; meanwhile, CSBV infection may attenuate honeybee immune responses by suppressing the expression of the key activator genes. Such as (i) down-regulated serine/threonine protein kinase and serine-type endopeptidase elaborately regulated the expression of serine proteinase and inhibit melanization via conversion of prophenoloxidase (PPO) to phenoloxidase (PO); (ii) down-regulated acyl-CoA Delta 11-desaturase and long chain fatty acids protein, thus disturb the fatty acid biosynthesis and immune. Our study offers important insights into understanding the mechanism of pathogenicity of CSBV and may lead to new molecular tools for both viral diagnosis and control for the bee viral diseases, especially native bee species. But new studies will test whether specific serine proteases are really involved in the melanization signaling pathway to protect honeybees against virus, as well as the immune potential of fatty acid biosynthesis.

## MATERIALS AND METHODS

### Sample collection

The samples were collected from Liaoning and Guangdong provinces in October 2016. The larvae of *A. cerana* were collected from 3 apiaries in different regions (Longmen, Meizhou, and Conghua) in Guangdong Province in China. CSBV-infected larvae were taken from the naturally infected colonies with obvious cystic phenotype symptoms in Liaoning and Guangdong provinces. The appearance of CSBV was proved according to the typical symptoms and reverse transcription polymerase chain reaction (RT-PCR) according to the protocols of Ma et al. **(7)**. The normal control larvae were from healthy larvae collected from three different apiaries (Huludao in Liaoning Province, Benxi in Liaoning Province, and Guangzhou in Guangdong Province). For RNA sequencing samples, the larvae of *A. cerana* were mixed into three groups. To avoid interferon by other viruses, PCR was performed to identify whether they were infected by the other 6 common honeybee viruses before RNA sequencing **(Table S10)**.

The 2^nd^ and 3^rd^ instar larvae were used for analysis since they were most susceptible to CSBV **(2)**. The age of larvae was identified by confining a queen to lay the eggs within 24 h. Twenty 2^nd^ larvae were considered as one treatment group, and each group consisted of three replicates. However, only two group samples were used for further analysis due to the RNA-seq data quality. These samples alive were taken and then immediately transferred into liquid nitrogen until use.

In addition, as follow-up supplementary experimentas material, fifteen CSBV-naturally-infected and healthy 4^th^, 5^th^, 6^th^ and 7^th^ larvae were used considered as one treatment group with three replicates, respectively. The larvae of *A. cerana* were collected from 6 different colonies in Guangdong Province Meizhou in October 2018, respectively.

### RNA extraction, library construction, and RNA-Seq

Total RNA of each sample were isolated from the larvae using the Trizol Reagent (Life technologies, California, USA). RNA quality was measured using an Agilent 2100 Bioanalyzer (Agilent Technologies, Inc., Santa Clara, CA, USA). The mRNA was obtained by NEBNext Poly (A) mRNA Magnetic Isolation Module (NEB, Ipswich, MA, USA). Briefly, the enriched mRNA was fragmented into approximately 200-nt RNA inserts, which were used to synthesize the cDNA. The end-repair/dA-tail and adaptor ligation were performed to the double-stranded cDNA. The corresponding fragments were obtained by Agencourt AMPure XP beads (Beckman Coulter, Inc. USA) and PCR amplification. PCR was performed by using Phusion High-Fidelity DNA polymerase, Universal PCR primers and Index (X) Primer. PCR products then were purified (AMPure XP system) and library quality was assessed on the Agilent Bioanalyzer 2100 system. At last, the clustering of the index-coded samples was performed on a cBot Cluster Generation System using TruSeq PE Cluster Kit v4-cBot-HS (Illumina, Inc. USA) according to the manufacturer’s instructions and the library preparations were sequenced on an Illumina Hiseq 2500 platform and paired-end reads were generated.

For sRNA library construction, we used the Small RNA Sample Prep Kit (Illumina) and 5 µg of RNA as the starting amount of the sample, and then 6 µL of water was filled up in total volume. Since the small RNA has a phosphate and hydroxyl group at the 5’ and 3’ end, respectively, the T4 RNA ligase 1 and ligase 2 (truncated) were respectively ligated to corresponding ends of the small RNA. After ligation, the ligated RNA fragments were reverse transcribed using M-MLV reverse transcriptase (Invitrogen, Inc. USA) and then the resultant cDNA products were amplified with two primers corresponding to the ends of the adapter sequences. Polyacrylamide gel was used to get small RNA libraries by electrophoresis and rubber cutting recycling. Finally, to perform clustering and sequencing of sRNA, cluster generation was performed by using the TruSeq PE Cluster Kit v4-cBot-HS (Illumina, Inc. USA) according to the manufacturer’s protocols. After cluster generation, the library construction was loaded to an Illumina Hiseq 2500 platform and sequencing paired-end reads were created.

### RNA sequencing quality control

Raw data with FASTQ format were analyzed through in-house perl scripts. Clean reads were obtained by removing those reads containing adapter, ploy-N, and low-quality. Reads with smaller than 18 nt or longer than 30 nt were trimmed and cleaned. Meanwhile, several parameters of the clean data were calculated such as Q20, Q30, GC-content and sequence duplication level. Low-quality reads, such as only adaptor, unknown nucleotides > 5%, or Q20 < 20% (percentage of sequences with sequencing error rates <1%), were removed by perl script. All the downstream analyses were based on clean data with high quality. The clean reads filtered from the raw reads were mapped to honeybees (*Apis mellifera*) (Aml-4.5) **(69)** and *A. cerana* genome **(39)** using Tophat2 software (Kim et al., 2013). The aligned records from the aligners in BAM/SAM format were further examined to remove potential duplicate molecules. Gene abundance were calculated based on the value of the TPM (transcripts per million).

### Identification of differential gene expression and sequence annotation

DESeq **(70)** and Q-value were employed to evaluate differential gene expression between CSBV and control groups. Gene expression levels were estimated using FPKM values (fragments per kilobase of exon per million fragments mapped) using Cufflinks software **(71)**. The false discovery rate (FDR) control method was used to identify the threshold of the P-value in multiple tests to compute the significance of the differences. Here, only genes with an absolute value of log2 Foldchange ≥1.5 and FDR significance score <0.05 were used for subsequent analysis. Genes were compared against various protein databases by BLASTX, including the National Center for Biotechnology Information (NCBI) non-redundant protein (Nr) database, and Swiss-Prot database with a cut-off E-value of 10^−5^. Furthermore, genes were searched against the NCBI non-redundant nucleotide sequence (Nt) database using BLASTn by a cut-off E-value of 10^−5^. Genes were retrieved based on the best BLAST hit (highest score) along with their protein functional annotation. Differentially expressed honeybee genes were analyzed against a background set of genes, which are all the *Drosophila* orthologues (Drosophila genome v. Dmel r5.42) in the honeybee genome (Amel4.5). The hierarchical clustering heatmap analysis of expression level of all genes was performed by MeV (https://sourceforge.net/projects/mev-tm4/).

### Identification of miRNA-targeted genes

Here, the sequences of ribosomal RNA (rRNA), transfer RNA (tRNA), small nuclear RNA (snRNA), small nucleolar RNA (snoRNA), and other kinds of non-coding RNAs were identified using a basic local alignment search tool (Bowtie tools soft, Version 1.1.2) against known non-coding RNAs deposited in the Silva database, GtRNAdb database, Rfam database and Repbase database, respectively, and unannotated reads of sRNA were obtained **(72)**. MiRDeep2 software (Version 2.0.5) was mainly used for the prediction of animal miRNAs **(73)**. Next, we mapped sRNA reads with lengths of 18-30 nt to the *A. cerana* genome **(69)** and CSBV genome (GenBank: KU574662.1) to get miRNA and CSBV-specific vsiRNA by using miRDeep2. Mapped reads were used to identify known miRNAs and novel miRNAs using miRBase 20.0 **(74)** and miREvo **(75)**. Using miRDeep2, based on the distribution information of reads on the precursor sequence (miRNA production characteristics, mature, star, loop) and energy information of precursor structure (RNAfold randfold), the Bayesian model was used to score and authenticate the miRNAs **(76)**.We then classified these small RNAs into several different categories according to their annotations.

Last, we used a well-established miRNA-target database MiRanda (v3.3a) and TargetScan (Version:7.0) to predict the target genes of the identified miRNAs and vsiRNA, and investigat the possible functions using the method of gene function annotation **(77)**. Target gene function was annotated based on the following databases: Nr (NCBI non-redundant protein sequences); Nt (NCBI non-redundant nucleotide sequences); Pfam (Protein family); KOG/COG (Clusters of Orthologous Groups of proteins); Swiss-Prot (A manually annotated and reviewed protein sequence database); KO (KEGG Ortholog database); GO (GeneOntology). The analysis of miRNA-mediated mRNA were performed based on the predicted miRNA-mRNA relationships. We selected miRNA-mRNA pairs showing opposite expression changes. The absolute value of log2 Foldchange ≥ 1 was used as a cutoff.

### GO and KEGG analysis

Unigenes were annotated in the NCBI non-redundant hits with gene ontology (GO) **(78)**. Perl script was then used to plot GO functional classification for the unigenes with a GO term hit to view the distribution of gene functions. The obtained annotation was enriched and refined using TopGo (R package). The gene sequences were also aligned to the COG database to predict and classify functions **(79)**. KEGG pathways were assigned to the assembled sequences by perl script. The interaction networks of targeted genes were predicted using STRING (http://string.embl.de/).

### Verification of expression difference genes by qPCR

The primers for the target genes are shown in **Table S11**. Approximately 1,000 ng of RNA was reverse transcribed using the PrimeScript RT Reagent Kit with gDNA Eraser (TaKaRa, Dalian, China) according to the manufacturer’s instructions, and the product was used as the template for qPCR. The qPCR reaction system consisted of 12.5 μL of 2 × SYBR Premix Ex TaqTM II (Takara, Dalian, China), 0.5 μL of upstream and downstream primers (10 mM), respectively, 2 μL of template cDNA, and 9.5 μL of double-distilled H_2_O in a total volume of 25 μL. qPCR was performed using a cycling sequence of 95 °C for 30 s, followed by 40 cycles of 95 °C for 5 s, 54-56 °C for 25 s, and 72 °C for 25 s. To normalize the qPCR result, according to previously reported CSBV study **(18)**, amplification of housekeeping gene β-actin was performed for each sample in this study. To verity the reliability of the RNA-seq result, twenty 2^nd^ larvae were used as one treatment experimental material, two biological replicates for each independent sample, and three technical replicates were analyzed for each biological replicate. While five 4^th^, 5^th^, 6^th^ and 7^th^ larvae were used as one treatment experimental material, three biological replicates for each independent sample, and three technical replicates were analyzed for each biological replicate.

### RT-PCR analysis for miRNA

For the analysis of miRNA expression, cDNA synthesis was performed using the miRCURY LNA Universal cDNA Synthesis kit II (Exiqon, Woburn, MA), according to the manufacturer’s instructions and using specific reverse transcription primer (5’-<u>CTCAACTGGTGTCGTGGAGTCGGCAATTCAGTTGAG</u>CCCACCAA-3’). RT-PCR were performed as follows: 94°C for 2 min followed by 30 cycles of 94°C for 30 s, 56°C for 35 s and 72 °C for 20 s. Specific sequence targeted miRNA 5 ‘-<u>ACACTCCAGCTGGG</u>GCTGGGCCTTCTTATTT-3’ was used as the forward primer, and URP, 5’-TGGTGTCGTGGAGTCG-3’ was used as the reverse primer. Data normalization of miRNA was carried out using the U6 reference gene (F: 5’-CTCGCTTCGGCAGCACA-3’, R: 5’-AACGCTTCACGAATTTGCGT-3’) **(55)**.

### Statistical analysis

The limma’s algorithm was used to filter the differentially expressed mRNAs and sRNAs, according to the significant analysis and FDR analysis **(80)**. All data analysis meets the following two criteria: (1) log2 FoldChange ≥ 1.5 or P < 0.05; and (2) FDR < 0.05. The relative expression of the target genes in the infected and control groups were calculated with the 2 ^-▵▵Ct^ method. Statistical comparisons and pearson correlation coefficient analysis were performed using GraphPad Prism 7 (GraphPad Software Inc., San Diego, CA, USA). Multiple t-tests were used to compare the significance of differential expression of genes between CSBV and control groups. The asterisks in the figures indicate significant differences: *P < 0.05; **P < 0.01).

### Data availability

The datasets generated or analyzed during the current study are available from the public database, http://gsa.big.ac.cn/ (CRA002310).

## ACKNOWLEDGMENTS

We acknowledge the support of The National Natural Science Foundation of China (31572471), the Agricultural Science and Technology Innovation Program (CAAS-ASTIP-2019-IAR), the Fundamental Research Funds for CAAS (Y2020PT17) and Young Talent Program of CAAS, GDAS Special Project of Science and Technology Development (2018GDASCX-0107).

## AUTHOR CONTRIBUTIONS

CSH conceived and designed the experiments. YCD, HXZ, SS, SY, DHY, and SD sampled and performed the experiments. CSH and YCD analyzed the data. CSH and YCD wrote and revised the manuscript.

## SUPPLEMENTARY MATERIAL

**FIG S1** The correlation analysis (r value = 0.8, P value = 0.05) between qPCR data and RNA-seq data veritied the reliability of the result.

**FIG S2** Expression pattern analysis of PPO gene was performed after CSBV infection at different instars by qPCR. **P<0.01.

**FIG S3** Length distribution of sRNA. (A) The length distribution of known miRNA. (B) The length distribution of novel miRNA. (C) The length distribution of CSBV-specific siRNA (vsiRNA).

**FIG S4** Expression pattern analysis of genes related RNAi pathway based on FPKM value was exhibited.

**Table S1** The data statistics of mRNA sample sequencing.

**Table S2** The statistic data of mRNA sequence comparison results between sample sequencing data and reference genome.

**Table S3** The data statistics of sRNA sample sequencing.

**Table S4** The genes list and their annotations from identified mRNA between blank and CSBV infected sick larvae.

**Table S5** The list of target genes and their annotations from 203 differential expressed mRNA between blank and CSBV infected sick larvae.

**Table S6** 23 differential expressed miRNAs between blank and CSBV infected sick larvae.

**Table S7** The list of target genes of siRNAs from CSBV-infected sick larvae.

**Table S8** The annotations on target genes of siRNAs from CSBV-infected sick larvae.

**Table S9** The TPM value of identificated siRNAs from CSBV-infected sick larvae.

**Table S10** Primer pairs for PCR used for detection the presence of common honey bee viruses.

**Table S11** Primers used to qPCR for target genes of siRNA.

